# An Empirical Study of ML-based Phenotyping and Denoising for Improved Genomic Discovery

**DOI:** 10.1101/2022.11.17.516907

**Authors:** Bo Yuan, Farhad Hormozdiari, Cory Y. McLean, Justin Cosentino

## Abstract

Genome-wide association studies (GWAS) are used to identify genetic variants significantly correlated with a target disease or phenotype as a first step to detect potentially *causal* genes. The availability of high-dimensional biomedical data in population-scale biobanks has enabled novel machine-learning-based phenotyping approaches in which machine learning (ML) algorithms rapidly and accurately phenotype large cohorts with both genomic and clinical data, increasing the statistical power to detect variants associated with a given phenotype. While recent work has demonstrated that these methods can be extended to diseases for which only low quality medical-record-based labels are available, it is not possible to quantify changes in statistical power since the underlying ground-truth liability scores for the complex, polygenic diseases represented by these medical-record-based phenotypes is unknown. In this work, we aim to empirically study the robustness of ML-based phenotyping procedures to label noise by applying varying levels of random noise to vertical cup-to-disc ratio (VCDR), a quantitative feature of the optic nerve that is predictable from color fundus imagery and strongly influences glaucoma referral risk. We show that the ML-based phenotyping procedure recovers the underlying liability score across noise levels, significantly improving genetic discovery and PRS predictive power relative to noisy equivalents. Furthermore, initial denoising experiments show promising preliminary results, suggesting that improving such methods will yield additional gains.

## 1 Introduction

Genome-wide association studies (GWAS) are used to identify genetic variants correlated with a target disease or phenotype. Population-scale cohorts, such as the UK Biobank (UKB) [1] and Biobank Japan [2], present the opportunity to deeply study complex polygenic diseases [3] as their large sample sizes increase power to detect associated variants via GWAS. However, to maximize GWAS power, a cohort must also be accurately phenotyped. Traditional strategies for accurate phenotyping, such as manual expert labeling, are costly, subjective, and scale poorly to population cohorts. The availability of high-dimensional biomedical data in biobanks has enabled novel approaches in which machine learning (ML) algorithms rapidly and accurately phenotype large cohorts with both genomic and clinical data [4–7], remedying the manual phenotyping challenges mentioned above. While early ML-based phenotyping methods relied on high-quality labels to train the ML model, recent work demonstrated that disease liability scores produced by a model trained on noisy medical-record-based labels improved GWAS power despite poor label quality [8]. This suggests that ML-based phenotyping can be extended to a wider range of diseases for which only lower quality electronic health records, such as self-report, hospitalization codes, and general practitioner notes, are available. However, to this date no rigorous work has quantified the effect of such noise on ML-based phenotyping results.

ML-based phenotyping is particularly powerful for diseases that follow a liability threshold model [9] and for which the underlying liability is well predicted by the model. One example is glaucoma, for which the vertical cup-to-disc ratio (VCDR) is a quantitative endophenotype that strongly influences glaucoma referral risk [10]. VCDR can be accurately predicted from color fundus images [5, 6, 10]: VCDR predictions from Alipanahi et al. [5] were shown to be strongly correlated with a test set graded by 2–3 experts (*R* = 0.89, 95% CI = 0.88 - 0.90).

In this paper, we investigate the ability of ML-based phenotyping methods to denoise corrupted labels and propose a hybrid approach that helps correct for such labels. Specifically, we use the VCDR values predicted by the model of Alipanahi et al. in UKB as a ground truth liability score for glaucoma, apply varying levels of random noise to generate noisy label sets, and train a variety of models on the noisy label sets. We evaluate downstream performance of the models both for their ability to predict VCDR directly and for their influence on genomic discovery. To the best of our knowledge, this is the first empirical study of methods for ML-based phenotyping denoising. We show that while even a small amount of label noise significantly decreases downstream genetic discovery and polygenic risk score (PRS) performance, the standard ML-based phenotyping procedure not only recovers underlying genetic hits but also significantly improves PRS predictive power relative to both the ground-truth and noisy GWAS. Initial explorations of integrated denoising approaches show promising preliminary results, further motivating this line of research.

## 2 Methods

### Experimental setup

In using the VCDR values predicted by the model of Alipanahi et al. [5] as the true underlying liability score for glaucoma, we assume that the corresponding GWAS and downstream genomic analyses run on these labels, hereafter *clean VCDR*, capture the true causal variants associated with glaucoma. We use these results as an upper bound in our denoising experiments. We choose to directly use this continuous representation to maximize the performance of our corrupted baselines since a noisy liability score will have improved power relative to some binarized equivalent [9]. This allows us to ignore the impact of the ML-based phenotyping procedure converting binary labels to continuous risk scores. We then generate various noisy label sets, hereafter *noisy VCDR*, by applying a multiplicative Gaussian noise vector to the clean VCDR labels for ϵ ∈ {0.1, 0.3, 0.8} and clamping to non-negative values:

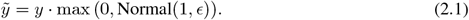

Results for additive random noise at similar levels had less impact on both ML performance and genomic discovery and are omitted from this manuscript. We use these noisy VCDR label sets in downstream analyses by both running GWAS directly on the noisy VCDR phenotypes as well as using the noisy VCDR values as targets to train ML models. The GWAS run on noisy VCDR serve as a lower bounds for our genomic analyses.

### ML-based phenotyping procedure

The ML-based phenotyping procedure consists of two separate processes: a model training phase (Figure 1a–c) in which we train an InceptionV3 network [11] to predict a patient’s target VCDR value from fundus imagery, and a model application phase (Figure 1d) in which model predictions are used for downstream genomic discovery in UKB. We describe standard networks trained using clean VCDR as *clean models* and standard networks trained using noisy VCDR as *noisy models*. Models that employ a modified training procedure to explicitly correct for noisy VCDR labels are referred to as *denoised models*. Each model is applied to UKB samples using a similar cross-fold strategy to prevent data leakage and the resulting liability scores are used to run GWAS. See Appendix A.1 for details.

**Figure 1:**
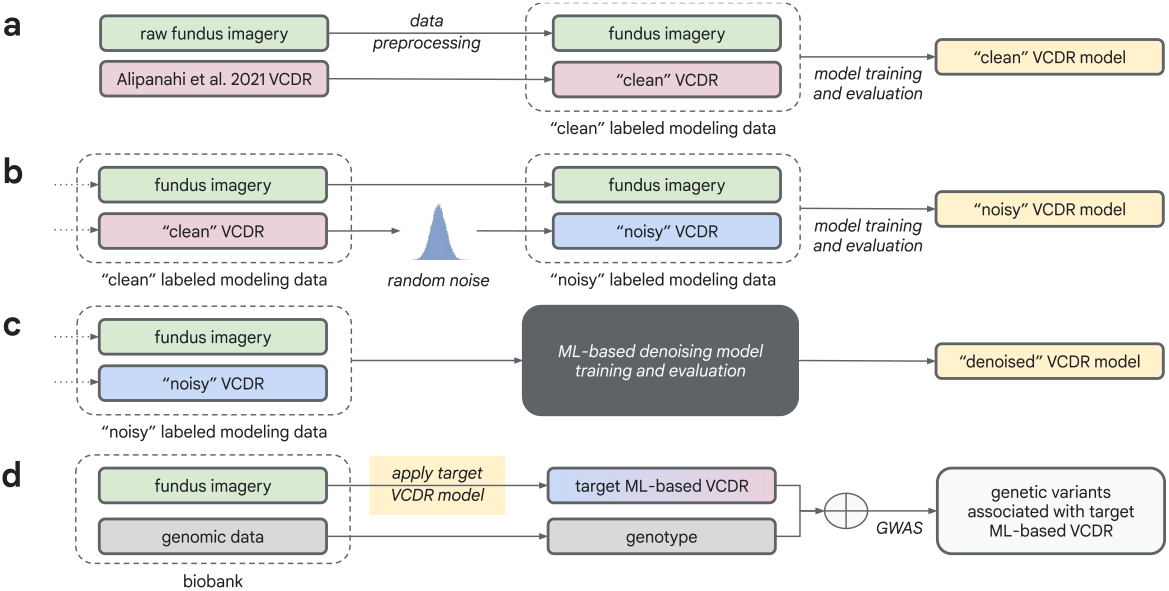
An overview of the ML-based phenotype denoising training and application processes for *clean, noisy*, and *denoised* VCDR models. Each model is applied to biobank fundus imagery and the resulting liability scores are used for genomic discovery.

### Performance evaluation

Though noisy models are optimized using their noisy VCDR targets, we evaluated all ML models according to their clean VCDR performance in the test set to assess each model’s ability to denoise the corrupted labels. We combined cross-fold predictions for each task to mimic the process used to generate GWAS phenotypes and evaluate the generalizability of the cross-fold methodology.

We evaluated improvements to genomic discovery by comparing noisy and denoised ML-based GWAS with our baseline clean and noisy label GWAS across three metrics: i) number of independent genome wide significant (GWS; *p* ≤ 5 × 10^-8^) hits [5, 12] from the clean VCDR GWAS replicated by ML-based GWAS, ii) statistical power of the GWAS, quantified using the non-centrality parameter (NCP) (Equation (2.2)), and iii) the predictive performance for clean VCDR of polygenic risk scores (PRS) summarizing the estimated effect of hits.

Running a GWAS is computationally expensive compared to the ML training process, often taking 10-20 times longer to complete. Since we have access to the underlying clean VCDR phenotype in our simulated setting, we were able to preemptively estimate ML-based GWAS power using NCP as a quantitative proxy. Equation (2.2) defines the NCP formulation for each variant *i*, where *β_i_* is the estimated effect size of *i*-th variant in the clean VCDR GWAS, se(*β_i_*) is the standard deviation of the estimated effect size in the clean VCDR GWAS, *m* is the total number of GWS variants for the clean VCDR GWAS, and *ρ* is the Pearson’s correlation coefficient between the target and clean VCDR phenotypes. The sum of per-variant NCP across all relevant hits, 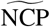, is then used as an indicator of expected GWAS power [13], which allows us to estimate downstream performance and to explore additional corruption and denoising methods with higher efficiency.

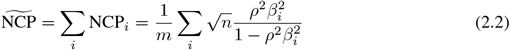

### Denoising attempts

After assessing the performance of standard ML-based phenotyping approaches under noisy conditions, we were interested in the application of denoising approaches to further improve robustness. There are many existing machine learning methods for reducing label noise or increasing training reliability [14–17]. For example, the Stratified Noisy Cross-Validation (SNVC) method [14] involves a two-step training procedure where one first trains multiple models on different folds of data and then uses the disagreement in prediction between these models to drop unreliable or hard-to-predict samples prior to performing a second round of training (Figure 5). Of candidate approaches, we considered SNVC the most suitable denoising approach. First, SNVC is agnostic to network architecture choice. Second, SNVC was designed for stratifying labeling noise in medical data and is expected to work on both binary and continuous labels, making it particularly compatible with our experimental setup.

We experimented with different types of filters for determining prediction disagreement. Distance-based filters computed L2 distance between predictions from different models and dropped samples using either a threshold of {1e-4,1e-3,1e-2} (*threshold filter*) or a percentile of top {10%, 20%} disagreement (*quantile filter*). Clinically binarized filters either binarized the prediction using clinically-defined thresholds of VCDR ≥ 0.7 [18] and considered the prediction reliable if the two models agree (*binary filter*) or expanded the 2-class binarization approach to 10-bin categorization (*multi-bin filter*). Lastly, to assess the upper bound of denoising performance, we designed an *oracle filter* where we took the ground truth levels of noise and dropped the top 20% of corrupted samples. Table 1 presents results for the two top performing filters: the clinically-defined binary filter and the oracle filter. Other filter approaches had similar or worse performance and the design of these filters deserves more study.

**Table 1:** ML and genomic discovery performance across labels and models. “ML Noisy *R*” and “ML Clean *R*” denote Pearson’s correlation between labels or model predictions and the target noisy or ground-truth labels, respectively. “NCP” denotes the non-centrality parameter as described in (2.2). “Replication %” captures the percent of ground-truth GWS hits replicated by the target phenotype’s GWAS. “PRS Euro *R*” and “PRS Non-Euro *R*” denote the Pearson’s correlation between PRS scores in the European holdout set (*n* = 1,472) and the non-European validation set (*n* = 10,095).

| Phenotype | ML Noisy $R$ | ML Clean $R$ | NCP | Replication % | PRS Euro $R$ | PRS Non-Euro $R$ |
| --- | --- | --- | --- | --- | --- | --- |
| Clean VCDR | - | $1.0000 \pm 0.0000$ | $194.7313 \pm 0.0000$ | 100% | $0.3047 \pm 0.0230$ | $0.2800 \pm 0.0096$ |
| Noisy VCDR ($\epsilon = 0.1$) | - | $0.9685 \pm 0.0005$ | $181.7726 \pm 0.0999$ | 92.86% | $0.1738 \pm 0.0239$ | $0.1509 \pm 0.0096$ |
| Noisy Model | $0.9511 \pm 0.0008$ | $0.9819 \pm 0.0004$ | $185.8213 \pm 0.0923$ | 96.75% | $0.3974 \pm 0.0223$ | $0.3652 \pm 0.0096$ |
| Denoised Model (binary) | $0.9506 \pm 0.0008$ | $0.9819 \pm 0.0003$ | $185.7674 \pm 0.0794$ | 96.10% | $0.4083 \pm 0.0221$ | $0.3712 \pm 0.0092$ |
| Denoised Model (oracle) | $0.9484 \pm 0.0008$ | $0.9795 \pm 0.0004$ | $184.8610 \pm 0.1024$ | 96.75% | $0.4008 \pm 0.0225$ | $0.3743 \pm 0.0094$ |
| Noisy VCDR ($\epsilon = 0.3$) | - | $0.7941 \pm 0.0035$ | $118.6192 \pm 0.4719$ | 76.30% | $0.1347 \pm 0.0240$ | $0.1188 \pm 0.0095$ |
| Noisy Model | $0.7677 \pm 0.0036$ | $0.9664 \pm 0.0006$ | $178.2096 \pm 0.1386$ | 93.51% | $0.3904 \pm 0.0222$ | $0.3539 \pm 0.0093$ |
| Denoised Model (binary) | $0.7669 \pm 0.0038$ | $0.9661 \pm 0.0007$ | $178.4808 \pm 0.1500$ | 94.16% | $0.3904 \pm 0.0231$ | $0.3607 \pm 0.0096$ |
| Denoised Model (oracle) | $0.7713 \pm 0.0037$ | $0.9729 \pm 0.0005$ | $181.6247 \pm 0.1259$ | 95.13% | $0.3933 \pm 0.0219$ | $0.3645 \pm 0.0092$ |
| Noisy VCDR ($\epsilon = 0.8$) | - | $0.4932 \pm 0.0070$ | $43.0540 \pm 0.5012$ | 41.88% | $0.0975 \pm 0.0261$ | $0.1451 \pm 0.0105$ |
| Noisy Model | $0.4652 \pm 0.0069$ | $0.9443 \pm 0.0009$ | $168.1538 \pm 0.2195$ | 91.56% | $0.4009 \pm 0.0217$ | $0.3596 \pm 0.0094$ |
| Denoised Model (binary) | $0.4610 \pm 0.0071$ | $0.9373 \pm 0.0011$ | $164.3056 \pm 0.2331$ | 89.94% | $0.3948 \pm 0.0232$ | $0.3719 \pm 0.0094$ |
| Denoised Model (oracle) | $0.4642 \pm 0.0071$ | $0.9463 \pm 0.0011$ | $168.9465 \pm 0.2057$ | 94.16% | $0.3845 \pm 0.0220$ | $0.3574 \pm 0.0096$ |

## 3 Results

We first evaluated the effect of artificial noise on GWAS power by comparing the clean labels and the noisy labels. We observed that the GWAS replication and PRS performance dropped as expected when the VCDR labels were corrupted, and that the power reduction is monotonic with the increased levels of noise introduced (ϵ = {0.0, 0.1, 0.3, 0.8}). We used these results as the baseline performance in the following results with ML-based phenotyping and denoising.

To estimate model performance under a real-world scenario with noisy medical measurements, we evaluated the change of the predictive power by comparing among the clean labels, the noisy labels, and the ML-based VCDR values using standard models trained with noisy labels, i.e., noisy VCDR models. Our results demonstrate that noisy models, though trained using noisy VCDR labels, have predictions that remain well correlated with the underlying clean VCDR values (*R* = 0.98,0.97, and 0.94, respectively). These results indicated the ML-based phenotyping methodology is robust against noise with respect to phenotype prediction (Figure 2c–d).

**Figure 2:**
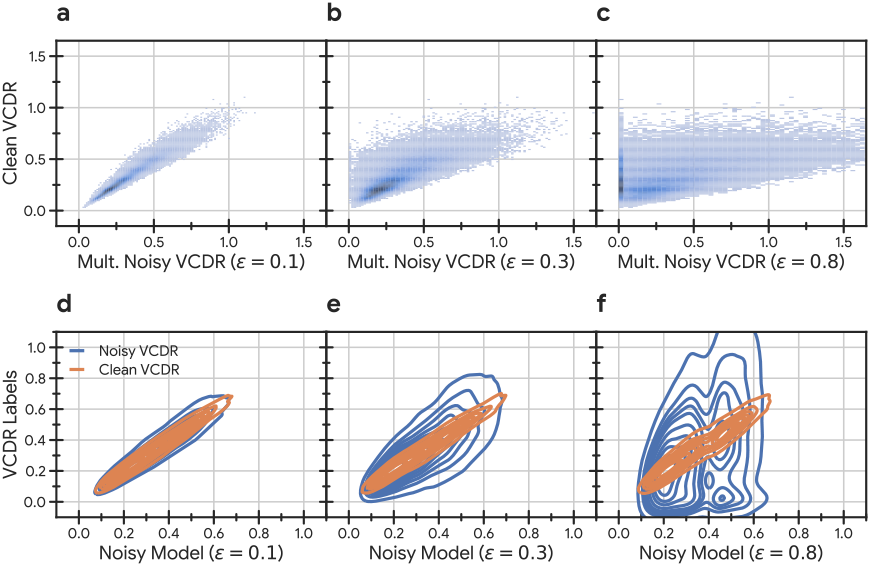
The robustness of ML-based phenotyping approaches to corrupted labels. a-c) Clean VCDR labels compared with their multiplicative noise equivalents. d-f) Noisy model predictions compared with the corresponding noisy labels and the underlying ground truth liability scores.

GWAS results were consistent with our observations of ML performance, showing that though the overall power was reduced when using noisy data, the reduction is much less when using ML-based VCDR values produced by noisy VCDR models trained to predict that same noisy data. We also observed that adding data corruption does not result in false positive genetic hits, which might indicate that the GWAS pipeline is robust against unbiased random noise or variance and therefore should be less affected by random data mislabeling or random errors in clinical measurements. Interestingly, noisy ML-based PRS models outperform not only their noisy label equivalents but also the ground-truth VCDR PRS.

In order to further improve model robustness, we added SNVC-based denoising methods on top of the original ML-based phenotyping model training procedure (Section 2). We aimed to empirically optimize the training phase, but also wanted to examine the theoretical upper bound of the SNVC method in denoising performance. Since, by experimental design, we had access to the ground-truth noise levels that were applied to each sample, we were also able to estimate the best possible denoising performance by designing an *oracle filter* that dropped samples according to those ground-truth levels. We used SNVC with this filter to train denoised VCDR models and ran GWAS on the resulting ML-based VCDR values. Our results show a marginal improvement in replication and PRS performance using this denoising approach when compared to the standard ML-based phenotyping process. However, other disagreement filters resulted in little to no improvement over the standard process, suggesting that there is more work to be done in designing these denoising approaches in the context of ML-based phenotyping.

## 4 Conclusion

In summary, we examined the robustness of the ML-based phenotyping procedure to label corruption and proposed an SNVC-based denoising method. By adding varying levels of simulated random noise to ground truth liability scores for glaucoma, we were able to empirically reason about both the effects of random phenotypic noise on downstream model performance as well as capture its impact on genomic discovery. Our results show that standard ML-based phenotyping approaches successfully recover underlying liability scores given corrupted labels and that our integrated denoising approach allows for modest additional gains under oracle disagreement conditions. These promising initial findings highlight the potential for extending ML-based phenotyping to diseases for which only low quality labels are currently available and motivate further denoising research. Exciting avenues for future work include extending this analysis to the binary label setting to better mirror the nature of the electronic health records often found in biobanks, evaluating the impact of additional noise distributions–including structured noise–to better understand how ML-based phenotyping handles systematic dataset bias, and further improving integrated denoising methods.

## Data and Code Availability

Genotypes and phenotypes are available for approved projects through the UK Biobank study (https://www.ukbiobank.ac.uk). This research has been conducted under Application Number 65275. Code and detailed instructions for InceptionV3 model training, prediction, and analysis are available at https://github.com/Google-Health/genomics-research/tree/main/ml-based-vcdr.

## Acknowledgments and Disclosure of Funding

We would like to thank Andrew Carroll, Babak Behsaz, Nick Furlotte, and Taedong Yun for helpful discussions and feedback throughout the course of this project.

This work was done while BY was an employee of Google LLC. The remaining authors are employees of Google LLC and own Alphabet stock as part of the standard compensation package. This study was funded by Google LLC.

## A Appendix

### A.1 InceptionV3 model training and application

Models were optimized to minimize MSE validation loss using Adam [19]. For each task, we performed a sweep over learning and regularization hyperparameters using the Vizier optimization service [20] (Table 2). Image preprocessing hyperparameters were held constant across runs (Table 3). In order to prevent overfitting, we employed an early stopping [21] patience of 5,000 steps and selected the checkpoint that resulted in the minimum validation loss. We implemented our InceptionV3 network using TensorFlow 2.0 [22] and each model instance was trained on a single NVIDIA Tesla V100 GPU using mixed floating-point precision [23]. Unlike Alipanahi et al. [5], we did not use a learning rate scheduler or model averaging. Models were initiated with the official `tf.keras.applications` weights pre-trained on ImageNet [24].

**Table 2:** Hyperparameters tuned using the Vizier optimization service [20]. The Vizier service employed a Gaussian process bandit optimization algorithm [29] to run a total of 70 trials with at most 35 trials running in parallel.

| Hyperparameter | Candidate Values |
| --- | --- |
| Learning rate | Sampled from $(1e-7, 1e-1)$ at unit log scale |
| Weight decay | Sampled from $(1e-7, 1e-3)$ at unit log scale |
| Dropout | $[0.0, 0.1, 0.2, 0.4, 0.6, 0.8]$ |
| Batch size | $[16, 32]$ |
| Adam $\beta_1$ | 0.9 |
| Adam $\beta_2$ | 0.999 |
| Adam $\epsilon$ | 0.1 |
| Adam amsgrad | FALSE |
| Max training steps | 250,000 |

**Table 3:**
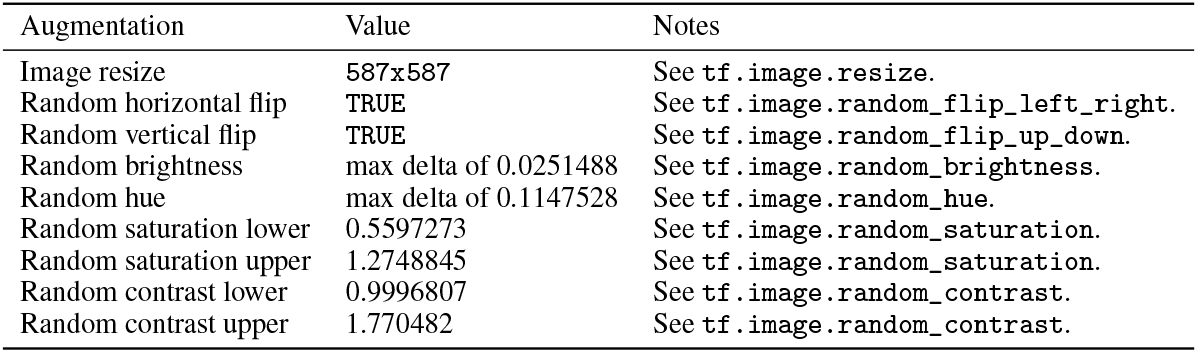
Image augmentation hyperparameters. These values were held constant across tasks and hyperparameter sweeps. Pixel values were first normalized to range [0, 1], augmented, and then clipped back to [0, 1]. Images were then centered to [−1, 1] as required by the InceptionV3 architecture.

To generate clean and noisy VCDR model liability scores, a separate model was trained on each fold and then applied directly to the other fold. Note that a separate hyperparameter search was completed for each fold. A dataset size ablation study and cross-fold model performance analysis showed strong pairwise correlations between a model trained on the full dataset (i.e., no cross-folding) and the two cross-fold models (Figure 4). Figure 5 illustrates the modified Stratified Noisy Cross-Validation (SNVC) [14] process used to generate denoised VCDR liability scores.

**Figure 3:**
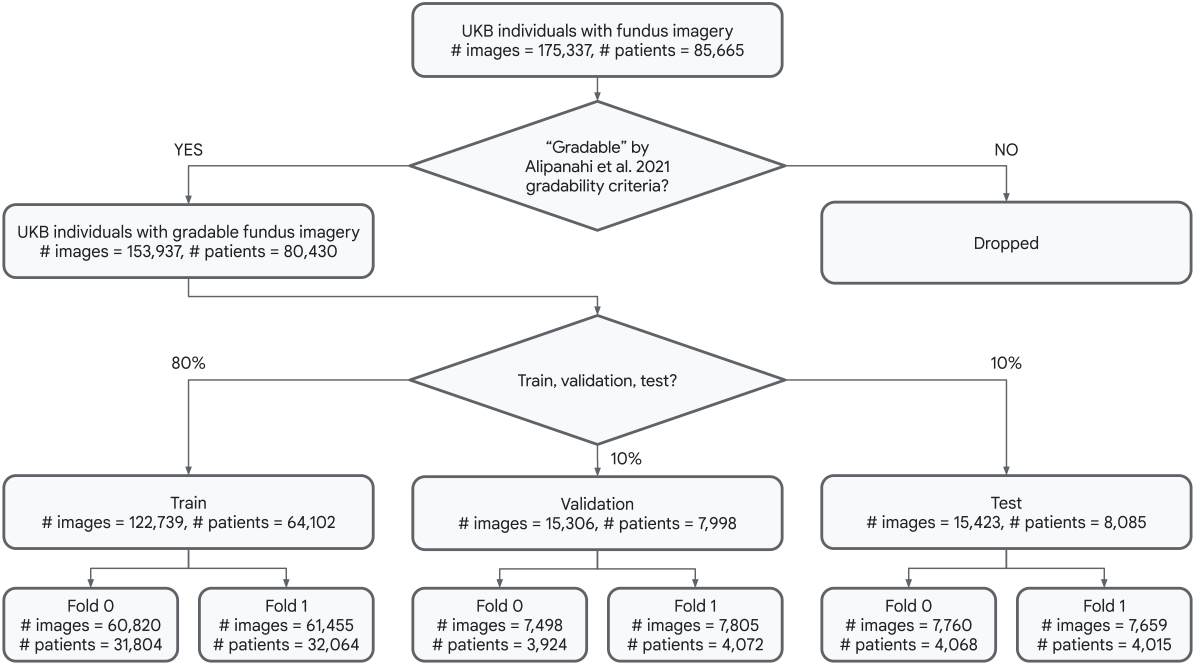
An overview of the ML train, validation, and test dataset splits and cross-folds. Individuals with gradable fundus images were randomly split according to their ID so that the split sizes are close to, but not exactly, the target train, validation, and test split distributions. Any individuals with estimated genetic relations spanning splits or folds were dropped (UKB fields 22011 and 22012).

**Figure 4:**
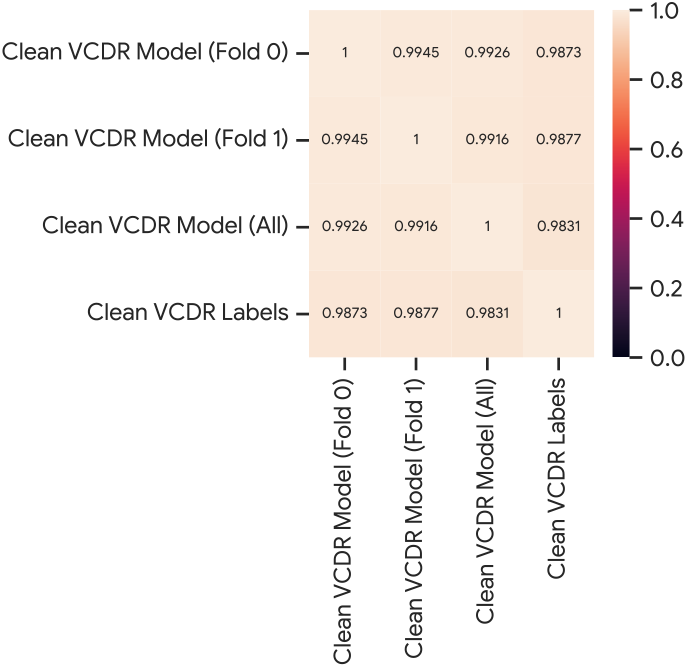
Pearson’s correlation (R) of validation set predictions across a model trained using only Fold 0 training samples, only Fold 1 training samples, and all training samples.

**Figure 5:**
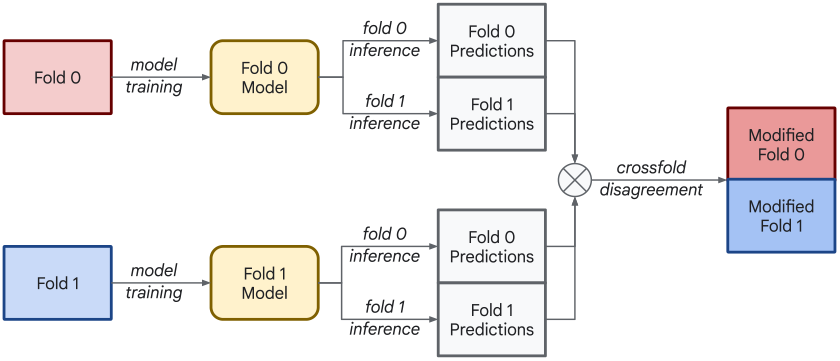
An overview of the modified Stratified Noisy Cross-Validation (SNVC) [14] process applied during the denoising process. Each model is trained on its respective fold and then applied to both folds. We calculated cross-fold model disagreement with a disagreement function and removed high disagreement individuals from the two folds. We retrained the models using their corresponding modified fold and then used each model to generate predictions for the other fold.

Models are trained on eye-level samples while GWAS is run on individual-level samples. We followed the eye-to-individual aggregation methods outlined in Alipanahi et al. [5], selecting images from only the first-available visit and then taking the mean of the target value across images.

### A.2 Genome-wide association studies and polygenic risk scores

GWAS analyses of all VCDR labels and ML-based VCDR phenotypes were performed using BOLT-LMM v2.3.6 [25, 26]. Prior to GWAS, all phenotypes were rank-based inverse normal transformed (INT) to increase the power for association discovery [27]. We controlled for sex, age at visit, visit ID (i.e., either 1 or 2 to indicate the first or second visit), number of eyes used to compute the target phenotype, average fundus image gradeability, refractive error, genotyping array, and the top 15 genetic principle components (PCs). GWS loci were obtained by merging hits within 250kb together. Polygenic risk scores (PRS) were computed using BOLT-LMM’s Best Linear Unbiased Prediction [25, 28].

